# Modeling collective cell migration in a heterogeneous environment: *Drosophila border* cell migration as a model paradigm

**DOI:** 10.1101/2025.10.16.682966

**Authors:** Soumyadipta Ray, Tuhin Roy, Sayan Acharjee, Gaurab Ghosh, Mohit Prasad, Dipjyoti Das

## Abstract

Collective cell migration underpins development and disease progression, yet the influence of heterogeneous microenvironments on intercellular forces and migration dynamics remains elusive. Focusing on the chemotactic migration of border cell clusters in *Drosophila* egg chambers, we present a computational framework that integrates intercellular force dynamics and cell shape changes during large-scale migration, validated with live imaging data. Our model replicates *in vivo* observations, such as rotations of the cell cluster and its tendency to follow a central path in the egg chamber. Using the model, we discover instantaneous acceleration/deceleration phases as the cell cluster migrates through junctions of other cell types. We also predict how environmental tissue topology and perturbations in chemoattractant gradients and adhesion impact cluster speed and rotation. By addressing the limitations of prior models of border cell migration, our approach offers broad applicability to other migration contexts involving interactions among different cell types in complex environments.

## INTRODUCTION

Collective cell migration is essential for various physiological processes, including embryonic development^1,2^, tissue homeostasis^3^, and cancer metastasis^4^. Cells can actively move by exchanging their neighbors through structural and shape changes, as seen in zebrafish tail elongation^5^, gastrulation invagination^6^, and wound healing^7^. Large-scale migration, covering distances much greater than cell size, often relies on chemoattractant or repellent gradients^7–13^. Examples include zebrafish posterior lateral line migration^14,15^, neural crest stream migration^16–18^, border cell movement in the *Drosophila* egg chamber^8,9,19,20^, and cancer progression^3,4,21^. These processes are influenced by guidance cues and interactions between cells and their microenvironment, composed of the extracellular matrix or other cells. While the effects of guidance cues are well-studied^9,19^, the impact of microenvironmental topology and heterogeneity on cell migration remains unclear.

Theoretical modeling has become an important complement to *in-vivo* live imaging for addressing key questions in developmental biology^9,19,22–28^. Two main modeling approaches are individual-based models (IBMs) and partial differential equation-based (PDE) models^29^. IBMs, particularly off-lattice frameworks representing cells as soft interacting particles, are more commonly used due to the challenges of incorporating cell-type heterogeneity in PDEs^10,29^. IBMs have been applied to various contexts of long-range cell migration, including Zebrafish tail elongation^1^, fish keratocyte movement^30^, neural crest migration^17,29^, Drosophila ventral furrow formation^31^, and Zebrafish lateral line formation^32^. However, they lack explicit modeling of cell shape changes and cell-environment interactions. In contrast, Vertex or Voronoi-based models^33–36^, which treat cells as extended objects rather than simple particles, effectively capture structural changes in cell collectives, such as rigidification of the zebrafish presomitic mesoderm^2^ and cell invagination during gastrulation^37^. Yet, these models primarily address neighbor exchanges in uniform cell populations without large-scale migration and are computationally intensive. This highlights the need for a framework that combines the strengths of particle-based and cell-shape-based models to simulate large-scale cell migration in heterogeneous environments.

Here, we focus on the collective migration of border cells in the *Drosophila* egg chamber, which consists of 15 large nurse cells surrounded by an epithelial layer^20^. During developmental stages 9–10, a cluster of 6– 10 follicle cells, known as border cells, migrates posteriorly from the anterior end towards the oocyte. As the cluster moves through relatively stationary nurse cell junctions, it carries two nonmotile polar cells^8,19^ (Fig. 1A, 1B, and movie S1). This migration is guided by two oocyte-secreted chemoattractant gradients: (i) an anterior-posterior gradient of PDGF- and VEGF-related factor 1 (PVF1) activating its receptor (PVR), and (ii) a mediolateral gradient of Spitz, Keren, and Gurken activating the epidermal growth factor receptor (EGFR). Blocking these receptors halts migration^8,19^. Border cells also maintain strong adhesion within the cluster to preserve integrity, while their adhesion to nurse cells is relatively weaker^38,39^. Intriguingly, the cluster exhibits two intertwined modes of dynamics: an anterior-posterior translational motion coupled with cluster rotation or tumbling, navigating along a central path of least resistance between nurse cells^8,40–42^. The underlying physical mechanisms driving these rich dynamics are still not fully understood.

**Figure 1.**
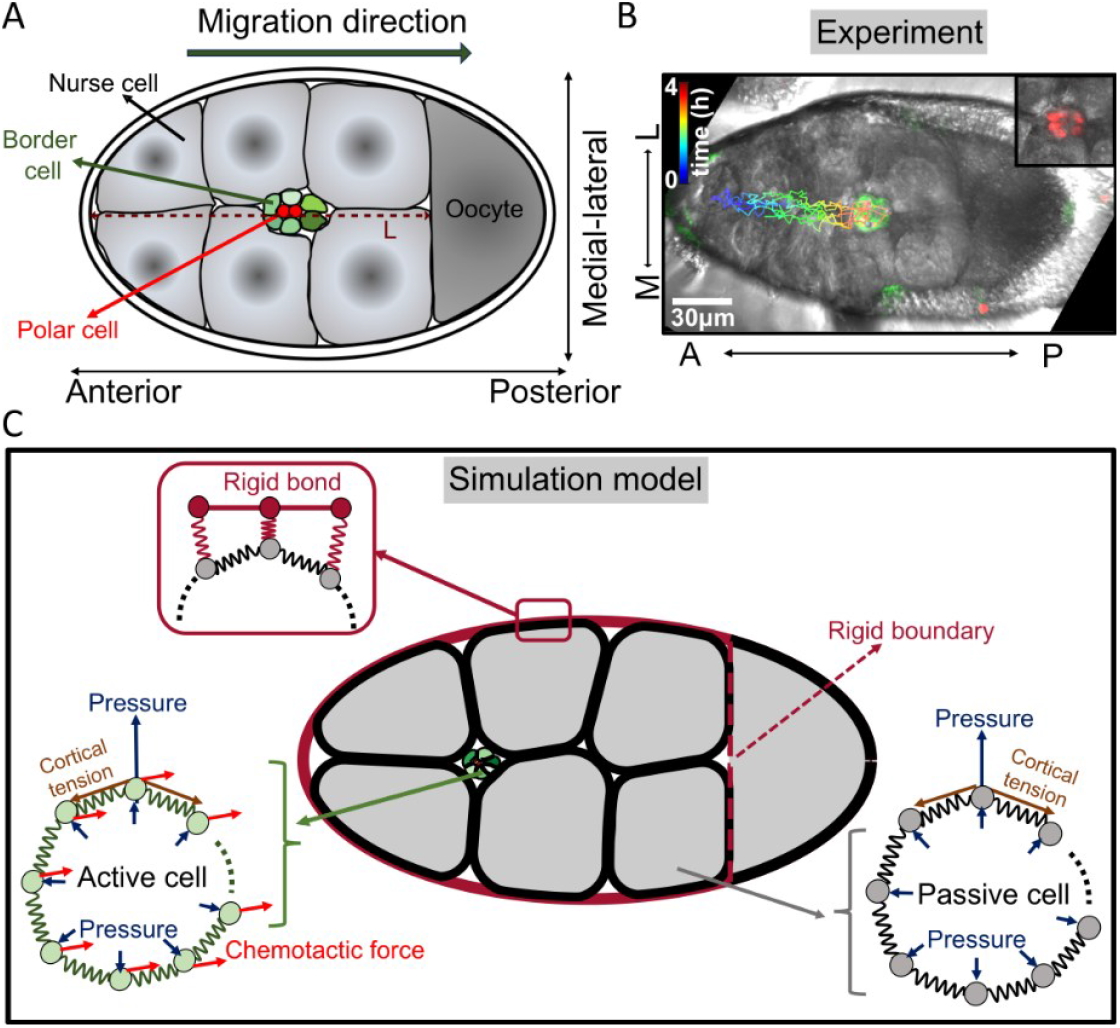
Simulation model of border cell migration in the *Drosophila* egg chamber. **(A)** A schematic of a *Drosophila* egg chamber illustrates the border cell cluster (green), carrying polar cells (red), migrating between nurse cells (light grey) toward the oocyte (dark grey). ‘L’ denotes the length of the egg chamber (brown dashed line). **(B)** A time-lapse snapshot of wild-type border cell migration shows cell membranes marked in green (by expressing *slbo*-Gal4,UAS-mCD8:GFP) and nuclei marked in red (UAS-*stinger-NLS*, inset). Images were captured using a confocal microscope with a DIC filter to identify nurse cell junctions. The tracked migration paths of border cell nuclei are overlaid, with a color gradient representing time. See Methods for experimental and analysis details. **(C)** Simulation framework: Each cell is modeled as a loop of beads connected by springs. Beads experience outward cytoplasmic pressure and tangential spring forces due to cortical tension (bottom-right). Border cells are assumed to be ‘active,’ i.e., their beads sense a chemotactic drag force (red arrows, bottom-left) combined with a directional noise towards the oocyte. The egg chamber’s outer boundary (brown) is assumed to be rigid, consisting of immovable particles that interact with nurse cell beads (top-left). See Methods for model details.

Existing theoretical and computational models offer crucial but partial insights into border cell migration. A theoretical study^22^ modeled chemoattractant-driven forces, treating the cluster as a spherical object but ignoring nurse cells. Similarly, Dai et al.^19^ simplified the cluster as a point particle moving through an effective potential, capturing its central path preference but neglecting detailed cell interactions. Another computational framework^40^ represented individual cells as aggregates of small spherical units with mechanical properties. However, these models overlook explicit interactions between heterogeneous cell types and fail to capture cell-level structural changes.

We present here a conceptually simple force-based model where individual cells are represented as soft, deformable objects (Fig. 1C). We explicitly model both nurse and border cells, capturing intercellular and chemotactic forces to account for the physical heterogeneities of the environment. Validated against live imaging data, the model reveals that the border cell cluster undergoes alternating phases of acceleration and deceleration driven by the topology of nurse cell junctions. Notably, we find that cluster rotations are more frequent near the junctions of three or more nurse cells than at two-cell junctions. Additionally, the model predicts the effects of disrupted chemoattractant cues on both translational and rotational motion. Perturbations to the relative strengths of border-border, nurse-border, and nurse-nurse adhesions significantly hinder migration, indicating the critical role of differential adhesion in this process. Overall, our framework enables precise control over cell-level mechanical parameters and intercellular interactions, offering insights into cell migration in heterogeneous environments.

## RESULTS

### Model

We combined features of cell shape-based^33–35,43–45^ and particle-based models^46–49^ to develop a 2D simulation framework for border cell migration. Building on deformable cell models^50–52^, we represent a single cell as a closed loop of beads connected by springs (Fig. 1C, bottom-left). Each bead experiences radially outward cytoplasmic pressure and tangential tension from adjacent springs, simulating actomyosin contractility. Beads from different cells interact via two short-range forces: attraction for cell-cell adhesion and repulsion to prevent interpenetration. Different cell types in the egg chamber are modeled by assigning distinct parameter values for intracellular pressure, spring tension, and intercellular adhesion (see Methods for details).

The egg chamber is modeled as an ellipse (Fig. 1C) enclosing six large nurse cells (since a 2D section along the anterior-posterior axis captures around six nurse cells). The posterior region (beyond 75% of the major axis from the anterior pole) represents the oocyte, separated from nurse cells by a rigid boundary (vertical dashed line in Fig. 1C). The chamber boundary is rigid, formed by immovable particles connected by stiff bonds, while maintaining attraction and repulsion with interior cells. At the simulation’s start, six border cells are placed near the anterior pole, forming a cohesive cluster with high mutual adhesion. A pair of polar cells are placed at the cluster’s center (Fig. 1). Nonmotile Nurse and polar cells are modeled as ‘passive,’ maintaining their shape through mechanical balance between pressure and cortical tension. In contrast, motile border cells are ‘active’ and experience chemotactic forces acting on beads exposed to the environment and interacting with nurse cell beads.

Border cells are guided by two chemotactic forces: one directed anterior-to-posterior (AP), driven by PVF1-PVR interactions, and another along the mediolateral (ML) axis, driven by the binding of Spitz, Keren, and Gurken ligands to EGF receptors. The concentrations of these oocyte-secreted ligands are assumed to follow exponential profiles^19,22^. Chemotactic forces are proportional to ligand concentrations but saturate at higher levels. Thus, the AP-directed force is modeled using a logistic function:

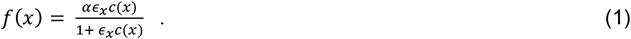

Here, *x* represents the x-component of the instantaneous bead position, *α* is a proportionality constant, and *ϵ*_*x*_ represents the ligand-receptor binding affinity. The ligand concentration is *c*(*x*) = *c*_1_exp (*x*/*ξ*_*x*_), where *ξ*_*x*_ is the length scale of the exponential decay. Similarly, the ML force is:

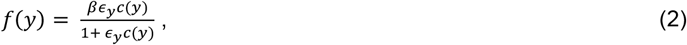

where *y* is the y-component of the bead position, *β* is the proportionality constant, and *ϵ*_*y*_ is the ligand-receptor binding affinity. The ligand concentration is *c*(*y*) = *c*_2_exp (*y*/*ξ*_*y*_). However, since the ML gradient is concentrated near the oocyte^19^, the ML-direction force is activated after the cluster has reached 2/3 of L, where L is the total migration length, measured as the distance from the anterior pole to the oocyte boundary^19^. Notably, the directions of chemotactic forces along the AP and ML axes are not fixed but include rotational noise, introducing variability in border cell polarity (see Methods). This accounts for the inherent heterogeneity in gradient sensing.

#### Unit Conversion

To make testable predictions, we scale our simulation to real units. The simulation time for complete AP migration under WT conditions is scaled to 5 hours, matching the average experimental migration time^19^. The migration length, L, is scaled to 150 µm, the typical egg chamber length^19,22^. For details on simulation procedures, see Methods.

### Cluster speed alternates between phases of acceleration and deceleration driven by nurse cell junctional topology

We compared predictions from wild-type (WT) simulations with *in vivo* live imaging data of border cell migration (simulation parameters in Table S1). In WT simulations, the border cell cluster migrated posteriorly through nurse cell junctions, reaching the oocyte while maintaining its architecture (Fig. 2A-C, Movie S2). Quantifying the net mean speed of the cluster center revealed that the speed was higher during the first half of migration than in the later half, consistent with previous experimental observations^8^ (Fig. 2D).

**Figure 2.**
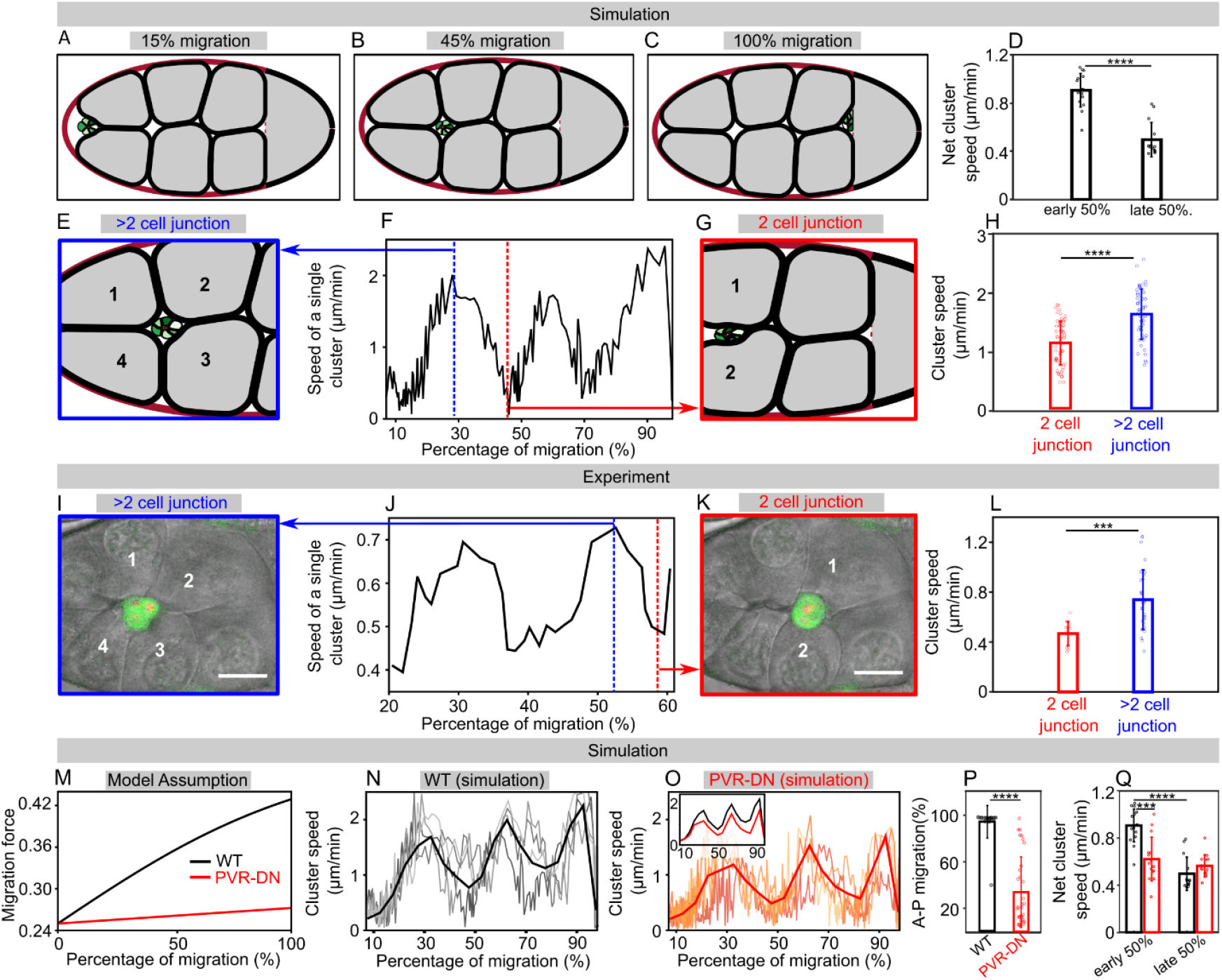
Border cell cluster alternates between acceleration and deceleration phases, influenced by nurse cell junction topology. **(A-C)** Snapshots from simulations show the anterior-posterior (AP) migration of border cells under wild-type (WT) conditions. **(D)** Net cluster speed during the early and late 50% of AP migration in WT simulations. **(E-H)** Instantaneous cluster speed exhibits oscillatory behavior, alternating between acceleration and deceleration phases. Sharp increases in speed correspond to 4-nurse cell junctions (blue dashed line in F, adjacent nurse cells are marked in E), while decreases in speed occur at 2-cell junctions (red dashed line in F, shown in G). Simulations predict significantly lower speeds at 2-cell junctions compared to junctions with more than 2 cells (H). **(I-L)** Experimental data confirm the model prediction. Oscillatory behavior in instantaneous speed aligns with the simulations, with high speeds at 4-nurse cell junctions (I, blue dashed line in J) and low speeds at 2-cell junctions (K, red dashed line in J). Instantaneous speeds measured from experimental movies also show lesser speed at 2-cell junctions than junctions between more than 2 cells (L). **(M)** Chemotactic migration forces in the model plotted with the distance from the anterior pole in WT and PVR-downregulated (PVR-DN) conditions. **(N-O)** Instantaneous speed profiles for WT (N) and PVR-DN (O) conditions. Individual simulation samples are shown as grey (N) and light red (O) curves, with average profiles in thick black (WT) and red (PVR-DN). The WT average speed is higher than PVR-DN (inset of O). **(P-Q)** AP migration percentages (P) and net cluster speeds in early and late AP migration (Q) for WT (black) and PVR-DN (red) simulations. Statistical significance (from t-test): ***P < 0.001, ****P < 0.0001. Experimental sample size: n=6 WT movies. Scale bars (I, K): 20 µm.

In simulations, we also measured instantaneous cluster speed by tracking the displacement of the cluster center between consecutive time points. We noticed oscillations in the instantaneous speed, alternating between acceleration and deceleration phases (Fig. 2F). We hypothesize that this behavior is influenced by nurse cell topology. Previous studies^19^ indicated that nurse cell organization biases the cluster toward a central path. We found that the cluster moves faster (blue dashed line, Fig. 2F) when passing through junctions of three or more nurse cells (Fig. 2E), while speed decreases significantly (red dashed line, Fig. 2F) at two-cell junctions (Fig. 2G). This suggests two-cell junctions produce greater resistance, significantly reducing the cluster speed (Fig. 2H).

To validate this prediction, we analyzed WT live imaging movies (Movie S1). Using UAS-Stinger-NLS to label border and polar cell nuclei (Fig. 1B), we tracked the cluster center while simultaneously visualizing nurse cell topology with a DIC filter. Experimental data confirmed alternating acceleration and deceleration phases in instantaneous speed (Fig. 2J). Faster speeds correlated with >2-nurse-cell junctions (Fig. 2I), while slower speeds corresponded to 2-cell junctions (Fig. 2K). Moreover, cluster speed was significantly lower near 2-cell junctions than >2-cell junctions (Fig. 2L), supporting model predictions.

We further explored the effects of perturbing chemoattractant cues on border cell migration. To simulate PVR downregulation (PVR-DN), we reduced the chemotactic force on border cells (Fig. 2M, details in Methods). Both WT and PVR-DN simulations showed oscillating instantaneous speed profiles in many simulations (Fig. 2N-O). However, the average instantaneous speed was consistently lower in PVR-DN (Fig. 2O, inset). Additionally, anterior-posterior (AP) migration was disrupted in PVR-DN, as previously reported in experiments^19^ (Fig. 2P). Net cluster speed during the first half of migration was also significantly reduced in PVR-DN, consistent with earlier *in vivo* measurements^8^ (Fig. 2Q).

It was shown that the decrease in overall cluster speed is associated with increased rotational cluster movement^8^. This suggests nurse cell junctional topology may also regulate rotational motion, which we explore in the next section.

### Surrounding nurse cell topology affects the rotational movement of border cell cluster

Tracking border cell nuclei in experimental movies revealed their positional rearrangements within the cluster. For example, a leading border cell at 50% migration shifted to a side position later at 64% migration, with another border cell assuming the leading position (Figs. 3A(I), 3B(I)). This switching coincided with the cluster moving from a 2-nurse cell junction (Fig. 3A(II)) to a 4-cell junction (Fig. 3B(II)), suggesting nurse cell organization influences cluster rotation.

**Figure 3.**
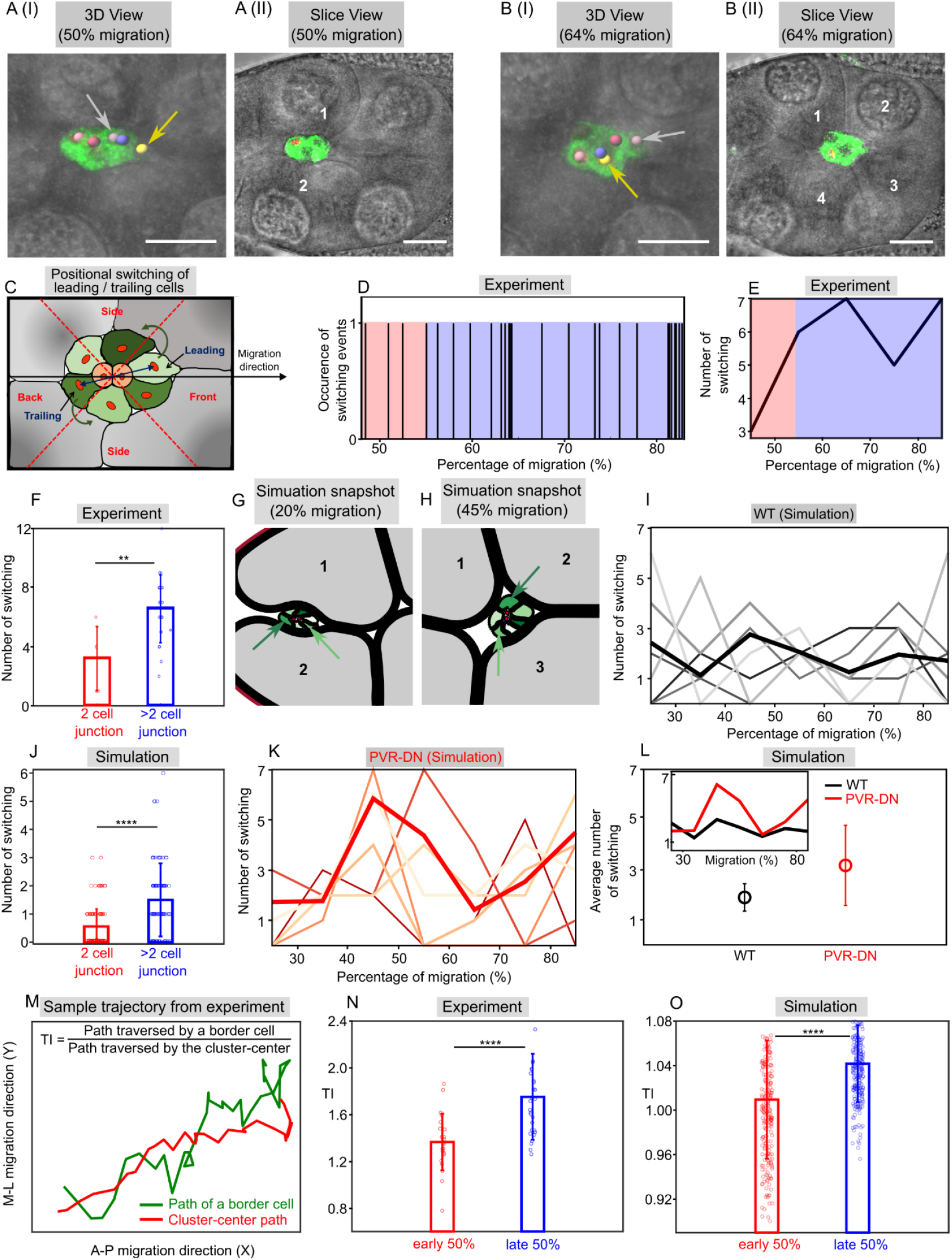
Nurse cell topology influences rotations of the border cell cluster during migration. **(A-B)** Images from a WT movie show positional rearrangements of border cells within the cluster as it migrates (colored balls mark border cell nuclei). Between 50% and 64% of AP migration, border cell nuclei switch positions (arrows in A(I) and B(I)). At 50% migration, the cluster passes through a 2-nurse cell junction (slice view in A(II)), while at 64%, it passes through a 4-cell junction (B(II)), highlighting increased positional switching near junctions with more than two cells. **(C)** Schematic of positional switching quantification: The cluster is divided into three zones (front, side, back) based on the angular position of nuclei relative to the migration direction. The leading border cell is closest to the direction of migration, and the trailing cell is furthest. A switching event occurs when the leading or trailing cell identity changes. **(D)** Events of leading or trailing cell switching over the migration percentage for the sample shown in A and B. The red and blue shaded regions represent times when the cluster passes through the 2-nurse cell and >2-nurse cell junctions, respectively. **(E)** The cumulative number of switching events (calculated from D) is higher near >2-cell junctions than 2-cell junctions. **(F)** WT experiments show significantly more switching events at >2-cell junctions compared to 2-cell junctions (from n=6 WT movies). **(G-H)** Simulation snapshots in WT condition at 20% (G) and 45% (H) AP migration show positional switching of border cells similar to experiments (indicated by arrows). **(I)** Cumulative number of switching events in WT simulations. Grey curves correspond to individual simulation samples; the bold-black curve denotes the sample average. **(J)** WT simulations show more border cell switching events near more than 2-nurse-cell junctions than 2-cell junctions (n=30 simulations). **(K)** In PVR-DN simulations, overall switching events increase throughout AP migration (Light-red curves denote individual simulations, and the bold-red curve is the sample average). **(L)** On average, PVR-DN simulations show more switching events than WT (L, inset: switching vs. migration percentage). **(M)** The cluster center’s tracked path (red) and an individual border cell’s path (green) from a WT movie indicate rotational behavior. The **tumbling index (TI)** quantifies rotation as the ratio of the individual cell’s path length to the cluster center’s path length. **(N-O)** TI is higher in the late 50% of migration compared to the early 50% in both WT experimental samples (N, n=6) and WT simulations (O, n=30). Error bars represent mean +/-std. P values are from the t-test (** P < 0.01, ***P < 0.001, ****P < 0.0001). Scale bars: 20µm.

To quantify these positional switchings, we divided the cluster into three zones—front, side, and back— based on the angle between the cluster’s instantaneous velocity and each border cell’s position relative to the cluster center (Fig. 3C). Border cells within 45° of this angle were in the front, those between 45°–135° in the side, and those beyond 135° in the back. The leading cell was then defined as the front cell closest to the cluster’s instantaneous velocity vector, while the trailing cell was the back cell furthest from it. A “positional switching” event was recorded when either the leading or trailing cell identity or both changed. We denote a single switching event and its absence as ‘1’ and ‘0’, respectively. Using this method, we tracked switching events throughout migration (Fig. 3D). Interestingly, switching events were more frequent when the cluster passed through >2-cell junctions (blue-shaded region, Fig. 3D) and less frequent at 2-cell junctions (red-shaded region, Fig. 3D). Aggregating the number of switching events over 10% migration intervals (from the same sample shown in Fig. 3D) showed significantly more switching at >2-cell junctions (Fig. 3E). Further, counting the total number of events from all experimental movies confirmed significantly higher switching near >2-nurse-cell junctions than 2-cell junctions (Fig. 3F).

Simulations also mirrored these observations. For example, in a sample simulation, a leading border cell moved to the side as the cluster passed a 3-nurse cell junction (Figs. 3G-H, Movie S3). Across multiple simulations, switching events alternated in frequency during migration, consistent with experimental data (compare Fig. 3I and 3E). Notably, switching events were significantly higher at >2-cell junctions than at 2-cell junctions, similar to our experimental observation (compare Fig. 3J and 3F). Our simulations also explain these observations: multi-nurse cell junctions, having larger intercellular spaces, offer less resistance to cluster motion than 2-cell junctions. As the cluster transitions from a 2-cell junction to a >2-cell junction, unbalanced interaction forces between nurse and border cells generate torque, altering angular momentum and inducing rotational motion. This rotation is also influenced by chemoattractant cues. In simulations of PVR-DN (reduced chemoattractant strength), switching events followed a similar pattern to WT (Fig. 3K) but occurred more frequently overall (Fig. 3L), consistent with prior experiments^8^. Thus, reduced AP gradient strength lowers cluster polarization but increases rotational motion.

To quantify rotation, we also measured the tumbling index (TI) used in previous experiments^8^. It is defined as the ratio of a border cell’s traveled path to the cluster center’s path (Fig. 3M). TI measurements from live imaging (Fig. 3N) and WT simulations (Fig. 3O) showed qualitative agreement, with higher TI values in the later 50% of migration, consistent with earlier reports^8^.

Since cluster migration depends on the intercellular spacing at junctions, it should also be influenced by differential adhesion between nurse and border cells, as adhesion regulates cell-cell gaps. In the next section, we thus explore the role of adhesion strength on border cell migration.

### Differential adhesion among border and nurse cells optimizes cluster migration

Using our computational model, we examined how the relative strengths of heterotypic adhesion (between nurse-border) and homotypic adhesion (between nurse-nurse and border-border) influence cluster migration. In the WT condition, complete AP migration occurred when border-border adhesion was significantly stronger than nurse-nurse or nurse-border adhesion (Fig. 4A; Table S1, Methods). This assumption, critical for maintaining cluster integrity, is consistent with earlier findings^38^. Increasing nurse-nurse adhesion to a level comparable to border-border adhesion severely disrupted AP migration (Fig. 4B, 4D). Similarly, increasing nurse-border adhesion strength comparable to border-border adhesion caused incomplete migration (Fig. 4C, 4D).

**Figure 4.**
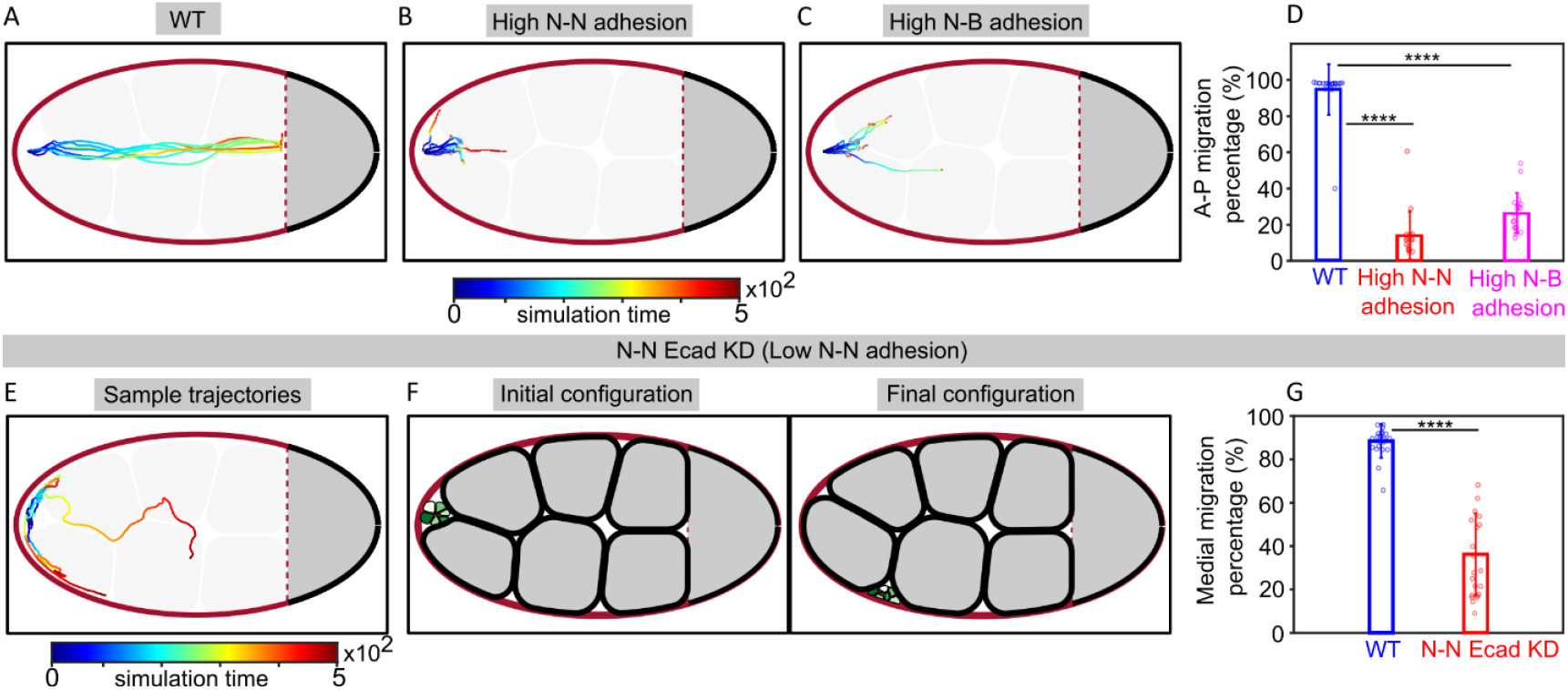
Differential adhesion between border and nurse cells is essential for efficient cluster migration. **(A-C)** Simulated migration trajectories for three conditions: WT (A), increased nurse-nurse adhesion (B), and increased nurse-border adhesion (C). **(D)** Both higher nurse-nurse (N-N) and nurse-border (N-B) adhesions significantly disrupt AP migration. **(E-F)** Reduced nurse-nurse adhesion, simulating the E-cadherin knockdown condition (N-N Ecad KD), results in disrupted medial migration. Simulation snapshots (F) show the border cell cluster taking a side path rather than a medial one. **(G)** The percentage of medial migration is significantly lower in N-N Ecad KD simulations compared to WT. The color gradient in trajectories represents simulation time. Statistical significance (from t-test): ****P < 0.0001.

Furthermore, previous experiments showed that knockdown of the Ecadh (Ecad KD) in the nurse-nurse junction causes defective medial migration^19^. Our model reproduced this migration defect. In simulations, we mimicked the Ecad KD condition by lowering the nurse-nurse adhesion and increasing the directional noise in chemoattractant forces (see methods). Directional noise likely arises from impaired anchoring of border cells to nurse cells^19,40^. Simulations under Ecad KD conditions showed disrupted medial migration, with the cluster moving sideways through the junction between nurse cells and egg chamber boundary, as observed experimentally (Fig. 4E, 4F; Movie S4). Medial migration was also significantly reduced in Ecad KD simulations compared to WT, aligning with the prior observation^19^.

## DISCUSSION

Border cell migration during oogenesis in the *Drosophila* egg chamber serves as a well-established model for studying collective cell migration in a heterogeneous environment. Here, we have developed a computational framework that integrates cellular forces, chemoattractant guidance cues, and individual cell shape dynamics during migration, validated by experimental observations. This model addresses a fundamental question: how do the microenvironment and cell-cell interactions influence the translational and rotational motion of the migrating border cell cluster.

Our simulations and experimental data analysis revealed that the border cell cluster alternates between acceleration and deceleration phases. The instantaneous cluster speed increases when passing through multi-nurse cell junctions and decreases when squeezing through two-nurse cell junctions. Cluster rotation and positional switching of border cells within the cluster are also reproduced in our model, as observed in live imaging^8,40^. These positional switching events are enhanced at multi-nurse cell junctions and suppressed at two-nurse cell junctions, aligning with our experimental data. This suggests that the topology of nurse cell junctions modulates cluster motion by generating variable resistance, which arises from the cluster encountering differing amounts of intercellular space during its movement.

Our model also shows that reducing posterior guidance cues (e.g., via PVR downregulation) hinders posterior migration and increases cluster rotation, consistent with prior experiments^8,19^. Additionally, perturbing the adhesion strengths among different cell types, such as border-nurse or nurse-nurse adhesion, substantially affects posterior migration. Notably, weakening nurse-nurse adhesion (mimicking nurse-nurse E-cadherin knockdown) disrupts medial migration, where the cluster cannot reach the oocyte and move sideways. These highlight the role of differential adhesion --- stronger border-border adhesion relative to nurse-nurse or nurse-border adhesion is essential for effective migration.

A key strength of our framework is its ability to integrate cell-level structural properties with force-based dynamics, enabling the simulation of large-scale migration with cell shape heterogeneity and neighbor exchanges. While polygon-based models (e.g., Vertex or Voronoi models) capture local shape changes and neighbor exchanges in processes like *Drosophila* wing disc formation or gastrulation^33,37^, they are computationally expensive for simulating heterogeneous environments with multiple cell types and large-scale migration. Notably, the border cell cluster in our simulation naturally follows a central path of least resistance without any imposed preference. This contrasts with the model by Dai et al.^19^, which treated the cluster as a single point particle in an ad hoc potential, ignoring cell-level details. Another theoretical study, which explored the relation between the mean speed and cluster size^22^, treated the cluster as a spherical object and overlooked the influence of the local microenvironment. Moreover, our model spontaneously generates cluster rotation even in 2D, driven by torque from imbalanced forces between the border and surrounding nurse cells. This is unlike a recent particle-based model^40^, which only captured cluster rotation in 3D but could not show cluster rotation in 2D. Our model thus removes some ad hoc features of earlier models, aligning more closely with experimentally observed phenomena.

Our framework offers broad applicability for investigating the mechanics of large-scale cell migration. We can generate experimentally testable predictions by precisely tuning the single-cell-level mechanical properties like cortical tension, cell-cell adhesion, and intracellular pressure. For instance, in zebrafish neural crest cell migration, lowering adhesion facilitates a fluid-like cluster state, promoting migration^17,53^ — such a behavior mimics solid-to-liquid phase transition in soft materials^54^, which our model can capture^50^. Another experiment on border cell migration reported a cytoplasmic pressure gradient in the nurse cells along the anterior-posterior direction^55^ ---this feature can be easily implemented by tuning the pressure magnitudes in our simulation. In the future, extending our model to 3D would enable the exploration of collective cell migration in biologically realistic geometries.

## RESOURCE AVAILABILITY

Custom-made codes for simulation (written in FORTRAN 90) are available in the GitHub Link: https://github.com/PhyBi/Border-cell-migration/tree/main. See the Key Resources Table in Supplementary Information for other details.

### Lead contact

Further information and requests for resources should be directed to and will be fulfilled by the lead contact, Dipjyoti Das.

### Materials availability

[Materials] generated in this study have been deposited in this paper.

### Data and code availability

1. Custom-made simulation code is available in GitHub. Link: https://github.com/PhyBi/Border-cell-migration/tree/main

## ACKNOWLEDGMENTS

DD acknowledges financial support from DBT and SERB, Govt. of India (DBT Project No. BT/RLF/Re-entry/51/2018 and ANRF project No. EEQ/2023/000551). DD and SR also thanks IISER Kolkata for the support.

## AUTHOR CONTRIBUTIONS

DD and MP conceived and supervised the project. DD supervised theoretical modelling and MP supervised experimental analysis. TR, SA, and GG designed and performed the experiments, analyzed the experimental data. SR developed the mathematical model, analyzed the data, and wrote the final manuscript with help from other authors.

## DECLARATION OF GENERATIVE AI AND AI-ASSISTED TECHNOLOGIES IN THE WRITING PROCESS

During the preparation of this work the author(s) used ‘Grammarly’ and ‘ChatGPT’ in order to improve the language and readability only in the Introduction and Discussion sections. After using this tool/service, the author(s) reviewed and edited the content as needed and take full responsibility for the content.

## Supplemental Information

### KEY RESOURCES TABLE

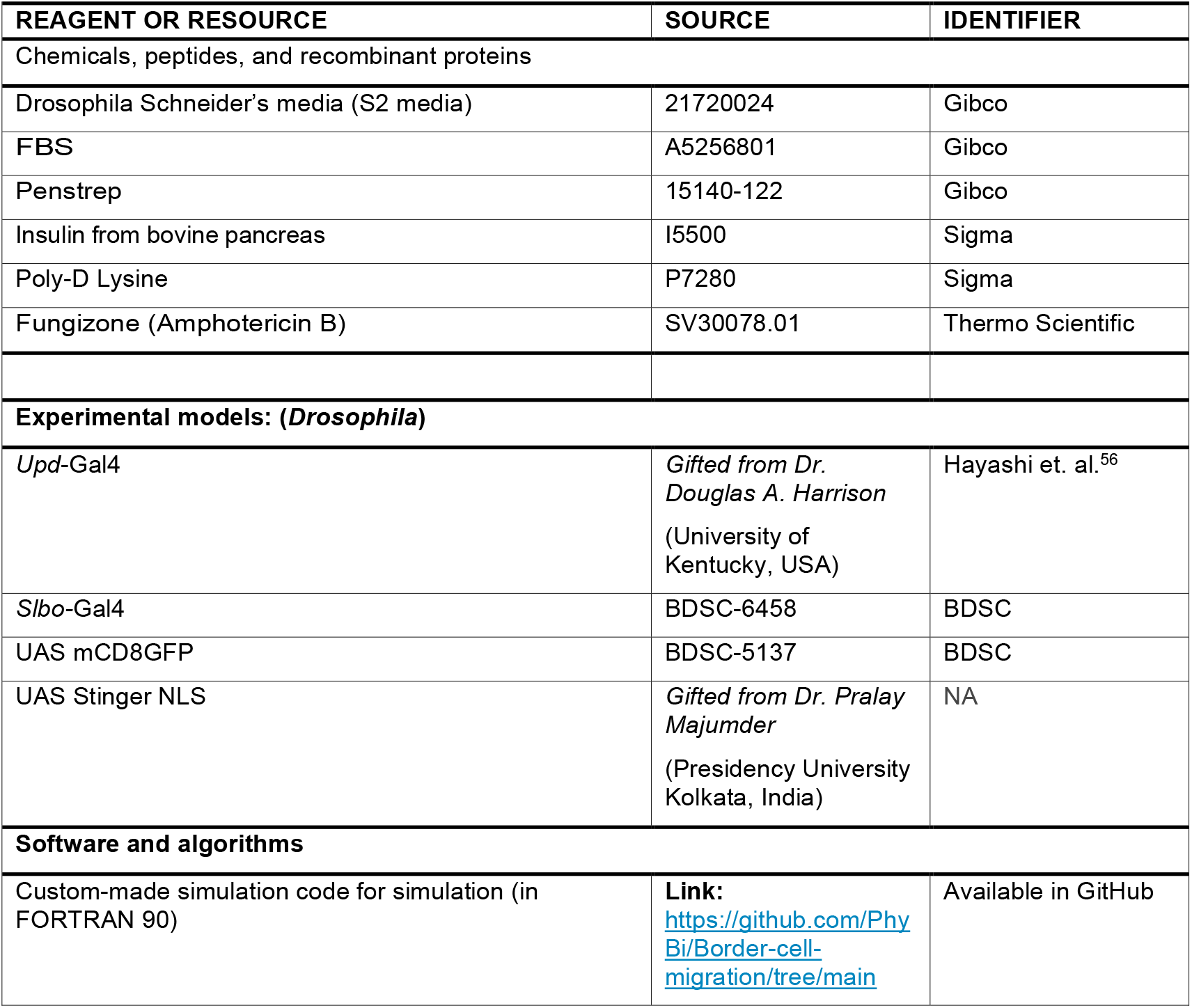

### EXPERIMENTAL MATHERIALS AND METHODS

#### Fly Strains

All stocks and crosses were maintained at 25°C. Experiments involving overexpression constructs of transgenes were fattened at 29 °C for 12 hours with yeast.

#### Setting the cross for live imaging

Setting the cross for live imaging: To mark the border cell and their respective nucleus, we crossed two fly stocks: *upd*-Gal4;*slbo-*Gal4,UAS-mCD8:GFP/Cyo with UAS-*Stinger-NLS*/Cyo and analyzed the progenies of interest. *Slbo*-Gal4,UAS-mCD8:GFP marks the border cell membrane while UAS-*stinger-NLS* labels all the nuclei of the migrating cluster.

#### Live imaging sample preparation

Non-curly flies from the cross (*upd-*Gal4;*slbo*-Gal4,UAS-mCD8:GFP/Cyo with UAS-*stinger-NLS*/Cyo) were collected. Around 5-6 female flies that were 2-3 days old were fattened in yeast-supplemented food media, with a few male flies at 29°C for 12 hours prior to the experiment.

Schneider’s *Drosophila* media (S2 media) supplemented with:

1. 15% FBS
2. 1% Penstrep
3. 0.1% Fungizone
4. 200 μg/mL Insulin.

For, our experiment 500 μL S2 media is prepared with 75 μL FBS and 10 μL insulin (10 mg/ml). The day before the live imaging, we had to prepare the Poly-D-Lysine coating in the coverslip. Poly-D-Lysine is referred to as PDL. This extracellular matrix, which is produced chemically, aids in cell adhesion to glass and plastic platforms used for tissue culture. For coating, we utilized 100 μg /ml PDL. With the aid of silica gel, we first adhere the coverslip to the confocal dish. Then, carefully pour 200 μL PDL of the aforementioned concentration into the coverslip. The microwave was then used to dry this until a suitable coating was evident. Coated coverslips are dried in 37°C incubator and UV treated for 15 minutes before use.

#### Imaging procedure

To perform the live imaging over the 4 hours span, we used the LEICA STELLARIS TCS-SP8 confocal microscopy. Confocal microscopy enables the visualization of specimens in three dimensions with excellent contrast and resolution. Confocal microscopy’s fundamental premise is to carefully scan a sample with a focused laser beam while detecting and collecting light only from one particular plane or depth inside the sample.

Even though the complete border cell migration takes around 5-6 hours, we performed the live imaging for 4 hours. Early stage 9 or late stage 9 egg chamber was selected, and images were taken at a 40x magnification for better spatial resolution. The image of the whole cluster was acquired at the green channel (488 nm) and the individual border cell nuclei were taken at the red channel (568nm). Following three to four trial runs to determine the optimal temporal resolution and Z width, a z width of 2 μm and a temporal resolution of 4 minutes were selected. Also, the following specifications like pixel size (1304 × 1304), and a scan speed of 550 Hz are maintained with bidirectional X mode with pinhole size -0.75 A.U. on during the image acquisition. In the initial part of the experiment, the DIC (Differential Interference contrast) was not set on while later we tend to acquire images with bright field or DIC filter along with the green and red channels. Additionally, for improved image capture in confocal microscopy, variables including zoom level, laser power, and gain power were adjusted. That is how a 3D video is generated where scans were done between the frames.

After imaging, the data was collected as a time series of z stack, meaning that at each time point, a set of images was built from various z planes. The imaging took place over the course of three to four hours at intervals of four minutes, so we have a minimum of 30-time frames with 12 to 15 z stacks each timeframe, depending on the z length, even if the z width is constant.

The output files were suitably labelled in the time and z planes and are of the LIF format for the TCS SP8 confocal and LOF format for the Stellaris confocal.

#### Visualization and quantification

The cells were tracked using IMARIS by BITPLANE, a program for 3D and 4D analysis in biological sciences. Using the IMS converter, these LIF and LOF files may be instantly converted to IMS (IMARIS file) format.

The spot-tracking feature builds a trail of the selected spots based on the spatial and temporal accuracy after we first choose a red channel and the average spot size (in our example, it is we consider 5 μm). An autoregressive motion algorithm was included in the software, which projected the location of the dots based on their continuous motion. Despite the fact that the analysis may be completely automated, a human and automatic tracking approach was used since certain frames needed to be manually altered due to a lack of spatial and temporal resolution. Before we could start spot tracking, we had to establish a standard reference frame for all of the videos. The DV and mediolateral axis cannot be distinguished from this imaging at this point, thus the AP axis of the egg chamber was assigned as the X-axis. Once the tracks were established, IMARIS reported the locations and speeds of the tracked points.

Border cells and polar cells are the two kinds of cells that make up the border cell cluster, as was previously mentioned. In the cluster’s core, polar cells were found, and following the net movement of the cluster and the polar cells showed that they are closely connected. The cluster’s center of mass (COM) is considered as the polar cells and cluster’s center was used to determine the trajectory of the COM. Using a large Spot with a diameter of around 20 μm, the green channel signal is tracked to complete the task. The IMARIS software does this auto-regressively. After tracking, each border cell nucleus’s position as well as the cluster’s coordinates’ center were stored in CSV format. Then we carry out further analyses using MATLAB.

### SIMULATION METHODS

#### MODEL

We model a single cell as a closed loop of beads connected by elastic springs with stiffness *K*_*s*_ and natural spring length *l*_0_ (see Fig. S1, Fig. 1C (main text)). Each bead experiences tangential tension forces due to the two adjacent neighbor springs, and a cytoplasmic pressure force, *Pl*_0_, directed outward-normal to the line tension of the given spring.

**Figure S1.**
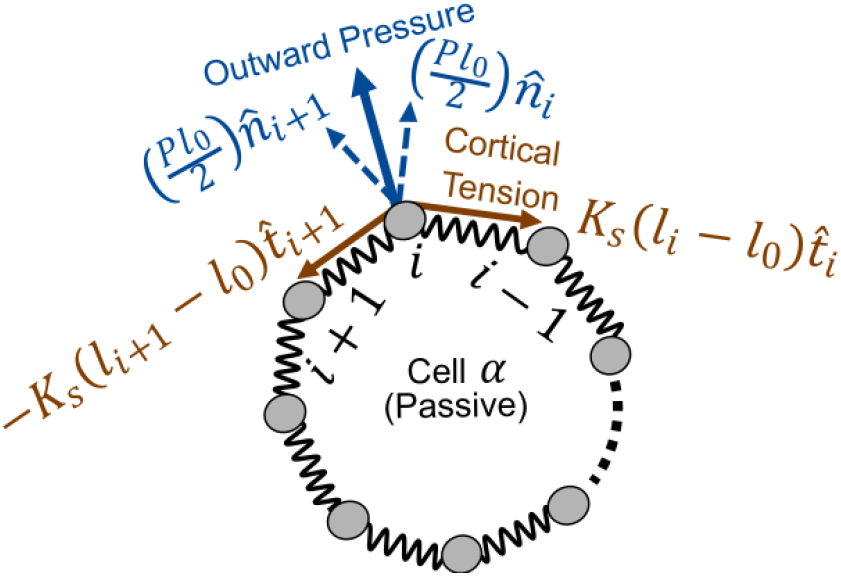
Bead-spring model for a single cell. The force components due to the tangential spring tension are shown in black arrows. The blue arrows are the normal components of the pressure force. The neighbors of the *i*-th bead are the (*i* − 1)-th and (*i* − 1)-th beads.

The intracellular force on the *i*-th bead is as follows:

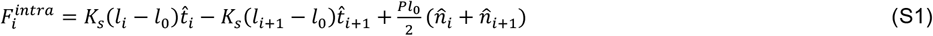

Here, *l*_*i*_ is the bond length between *i*-th and (*i* − 1)-th beads, and *l*_*i*+1_ is the bond length between (*i* + 1)-th and *i*-th beads, respectively. Also, 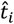 and 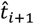 are the tangential unit vectors, and 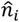 and 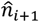 are the corresponding outward unit vectors perpendicular to 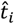 and 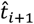, respectively (see Fig. S1). Now, for each specific cell-type, i.e., nurse, border, or polar cell (denoted by subscripts n, b, and p respectively), the force strengths naturally vary. We thus denote *K*_*s,n*_, *K*_*s,b*_, *K*_*s,p*_ as the spring strengths, *P*_*n*_, *P*_*b*_, *P*_*p*_ as the pressure force coefficients, and *l*_0,*n*_, *l*_0,*b*_, *l*_0,*p*_ as the natural spring lengths, and hence the intracellular forces as 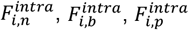 for the nurse, border, and polar cells respectively (for details, see table S1).

#### Cell-cell interactions

The intercellular force due to cell-cell interactions consists of two parts: (i) the Hookean attractive force, that approximates the cell-cell adhesion through adhesive proteins (like E-cadherins), and (ii) the Hookean repulsion that prevents the cell-cell interpenetration. Two beads of two distinct cells would come under the force of adhesion or repulsion only when the Euclidean distance between them is below the cut-off range of the attractive or repulsive forces. The interaction force on the *i*-th bead of *α*-th cell due to the *j*-th bead of *β*-th cell is as follows:

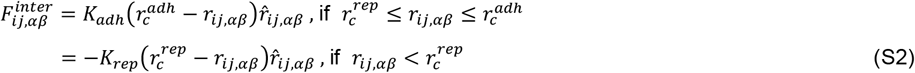

Where,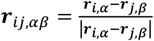 is the unit vector of the Euclidean distance between the two interacting beads. *K* _*adh*_ and 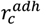 are the strength and cut-off range, respectively, for the attractive force, and *K* _*rep*_ and 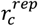 are those of the repulsive force. Now, the interaction strengths among two cells of same type or different types may differ. That’s why we have different strengths and interaction cut-off for adhesion and repulsion depending on the types of the interacting cell pair. See table S1 for details.

Now, if we consider all the contributions from other cells within the interaction ranges, the total force of interaction on the $i$-th bead of the $\alpha$-th cell would be:

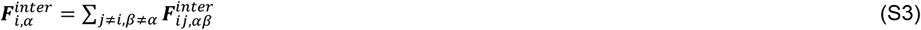

#### Active force term via the force of chemoattractant and Equations of motion

To model the chemoattractant force on the border cells, we incorporate a local, position dependent pulling force *f*(*x*) on the border cell beads that are exposed to the outer environment, making the border cells active. At a particular time instant, the border cell beads ‘exposed to the outer environment’ refer to the particular beads of a border cell, that do interact with adjacent nurse cells or have no interaction with other border cells or the polar cells. The chemoattractant forces act on the border cells in the anterior-posterior (AP) direction due to the binding of PVF1 ligands with PVR receptors, and in the medio-lateral (ML) direction (perpendicular to AP direction) due to the binding of Spitz, Keren, and Gurken ligands with EGF receptors. An exponential profile would be the most appropriate fit for the concentration of these oocyte secreted ligands. For low ligand concentration, the local pulling force is proportional to the concentration, while in high concentration, the pulling force is expected to reach a maximum value. Thus, we take the pulling force in AP direction as a logistic function of the chemoattractant concentration:

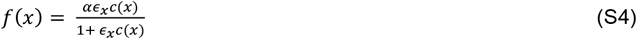

where *α* is a proportionality constant, and *ϵ*_*x*_ represents the ligand-receptor binding affinity. The ligand concentration profile is *c*(*x*) = *c*_1_exp (*x*/*ξ*_*x*_), where *ξ*_*x*_ is the length scale of the exponential profile decay. The variable *x* represents the x-component of the instantaneous position of the bead of a border cell. Similarly, the pulling force in the DV direction is:

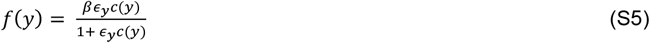

where *β* is the proportionality constant, and *ϵ*_*y*_ represents the corresponding ligand-receptor binding affinity. The ligand concentration is *c*(*y*) = *c*_2_exp (*y*/*ξ*_*y*_), where *ξ*_*y*_ is the length scale of the exponential decay. The variable *y* represents the y-component of the instantaneous position of the bead. The force *f*(*y*) in ML direction due to Gurken is activated only after the border cell cluster has reached 2/3 of the egg chamber length L.

#### Implementing the PVR-DN condition

We implement the PVR-DN scenario by increasing the decay length scale of the exponential gradient, *ξ*_*x*_ in the ligand concentration profile *c*(*x*), thus effectively reducing the local pulling force on the border cells (see table S1). It reduces the net pulling force on each of the border cells (see Fig. 2M in main text).

#### Active remodeling of the AP pulling force direction through noise

All the border cell beads ‘exposed to the outer environment’, get the pulling force in a specific direction at an instant. Then at a later instant, the direction gets remodeled due to the stochasticity in gradient sensing. We thus model the instantaneous AP pulling force direction as the rotationally diffusive cell polarity in self-propelled particle (SPP) model and deformable cell model. Thus, in vector notation, instantaneous direction of *f*(*x*) is 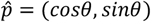 for all the border cell beads ‘exposed to the outer environment’. In the direction of 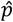, each bead possesses an AP pulling force of magnitude *f*(*x*). Although, the ML pulling force *f*(*y*) has been preserved solely towards the y-direction, keeping the model simple. Thus, at an instant, for the six border cells, all the border cell beads, those are ‘exposed to the outer environment’, have the same pulling force direction, but only the AP pulling force rotationally diffuses over time keeping the mean direction towards the AP-axis.

All together, the overdamped equation of motion for the *i*^*th*^ bead of a nurse cell (passive) is:

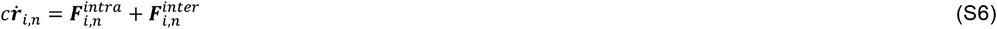

where, 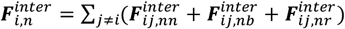

Here 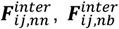, and 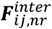 denote the nurse-nurse, nurse-border, and nurse-rigid boundary particles interaction forces respectively. As the polar cells are engulfed within the border cells, they don’t have any interaction with the nurse cells.

Similarly, the overdamped equation of motion for the *i*^*th*^ bead of a passive polar cell is:

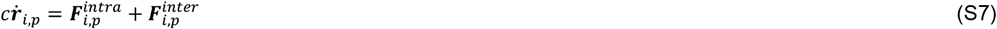

where, 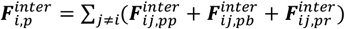

Here 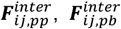 and 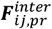 denote the polar-polar, polar-border, and polar-rigid boundary particles interaction forces respectively.

Finally, the overdamped equation of motion for the *i*^*th*^ bead of an active border cell is: 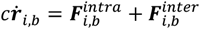, if the *i*^*th*^ bead is not ‘exposed to the outer environment’

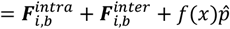, if *x*_*i,b*_ < 2*L*/3 and the *i*^*th*^ bead is ‘exposed to the outer environment’

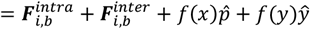, if *x*_*i,b*_ ≥ 2*L*/3 and the *i*^*th*^ bead is ‘exposed to the outer environment’

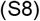

And, the AP pulling force direction 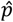, which is same for all the ‘exposed to the outer environment’ beads of the six border cells, also undergoes rotational diffusion over time, described as:

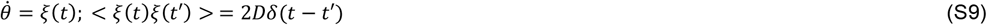

Where, *ξ*(*t*) is a Gaussian white noise with zero mean and variance 2*D*. The inverse of the variance sets the timescale of reorientation of the pulling force direction. The mean direction of the AP pulling force is towards the AP direction that has been chosen at the beginning of migration.

Here, 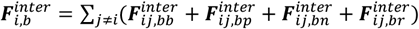

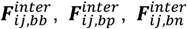, and 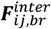 denote the border-border, border-polar, border-nurse, and border-rigid boundary particles interaction forces respectively.

### SIMULATION DETAILS

Here we consider *m*_*n*_ = 6 nurse cells in the two-dimensional model egg chamber along with *n*_*b*_ = 6 border and *n*_*p*_ = 2 polar cells consisting the cluster. Each of the nurse cells, border cell and polar cells consist of 100, 40, and 15 beads, respectively. The bead positions are updated by numerically integrating Equations S6-S9 using the Euler method. The time step is 10^−3^ and the total number of iterations is 5 × 10^5^.

#### Construction of the whole egg chamber architecture

##### Boundary of the egg chamber

We take an elliptic geometry for the egg chamber boundary with the major axis of the ellipse as the AP axis (along the x-axis in 2D) and the minor one as the ML axis (along the y-axis). Length of the major axis is taken twice the length of the minor axis as per experimental observation. We construct the rigid boundary by uniformly putting 1500 immovable beads along the perimeter of the ellipse (see the brown outer boundary in Fig. 1C, main text). Beads of other cells in the system interact through adhesion and repulsion with these immovable beads, maintaining the confinement of the cells within the chamber boundary. We keep the origin at the left-most point of the elliptic boundary, i.e. at the extreme anterior point. The portion of the egg chamber beyond 75% of the major axis length from the origin is modeled as the oocyte (at the posterior site), separated by a rigid boundary (vertical dashed line in Fig. 1C). The horizontal distance from the origin up to the oocyte boundary is denoted as the egg chamber length L (horizontal dashed line in Fig. 1A).

##### Constructing the nurse cells

We consider six nurse cells in our two-dimensional model egg chamber. The nurse cells are positioned within a specific region along the x-axis in the chamber, i.e., from the length of initial diameter of border cell cluster (discussed in the next sub-section) up to L, the rigid oocyte boundary (brown dashed line in Fig. 1C, main text). As in the live imaging data, we observe a positional symmetry of the nurse cells with respect to the AP axis, we also maintain the symmetry while seeding the nurse cell centers in simulation. Within the stipulated region along the x-axis, we first choose three of the nurse cell centers to be in the upper half of the chamber and rest three cells in the lower half, along with some randomness in their x and y components of position coordinates. The randomness is incorporated using uniformly distributed random numbers within a tolerable limit, so that the cell does not overlap with others and doesn’t cross the egg chamber boundary during the initialization. The initial diameter of the nurse cells has been taken as 20% of L as a rough estimate from experiments. We assign *n*_*n*_ = 100 beads for each nurse cell and choose the natural spring length as 0.1 simulation units which is of the same order of magnitude with the separation between successive beads in the egg chamber boundary, thus restricting the nurse cell beads to cross the boundary. The details of the parameters chosen for the nurse cells are in table S1.

##### Constructing the border cell cluster organization

We first roughly estimate the relative diameters of border cells, polar cells and the whole cluster with respect to the egg chamber length from experiments.

Accordingly, we choose the initial diameter of the whole border cell cluster as 13% of L and that of each border cell as 4% of L. The initial polar cell diameter is taken as half the border cell diameter. We then seed the cluster center at x = cluster radius and y = 0. The two polar cell centers are then positioned on the left and on the right side of the cluster center respectively, each having the distance of polar cell radius away from the cluster center along the x-axis. The six border cells centers are then arranged circularly in an anticlockwise manner, surrounding the two polar cells, taking center as the cluster center position. Then we assign *n*_*b*_ = 40 and *n*_*p*_ = 15 beads to each of the border and polar cells respectively. The natural spring lengths for border cells and polar cells are 0.048 and 0.046 simulation units respectively, and notably these are almost half of the natural spring length for the nurse cells. Note that the border-border and border-polar adhesions are much higher than other cell-cell adhesions in the model, to maintain the cluster cohesiveness and integrity. For detailed parameter values, see table S1.

#### Equilibrium tissue configuration through passive cell dynamics

The initially generated cell-system is then evolved by integrating Equation S6-S8 with *f*(*x*) = 0 and *f*(*y*) = 0, i.e., we evolve the cells passively. During this time window, the cells grow, and by interacting with each other, they approach a nearly space-filled equilibrium configuration. This equilibrium configuration then serves as an initial configuration for simulations of the active system. In all our simulations, we equilibrate the configurations up to the total iterations of 10^4^.

#### Simulation of the active system

After the passive system attained an equilibrium configuration (described above), we reset the time to zero. The border cells within the equilibrated tissue are then subjected to active migration forces as described in the previous section of ‘Active force’ and the corresponding border cell bead positions are updated through Equation S8 and S9. The other cell positions are updated with Equation S6-S7, as before. We evolve the system up to 5 × 10^5^ iterations by which the border cell cluster eventually reaches the oocyte boundary for the wild-type case. That’s why we chose 500 simulation time (5 × 10^5^ × *dt*) as the standard observation time window in-vivo as 5h. Across this 500 simulation time, we measure all the relevant observables to analyze the cluster dynamics.

##### Code availability

The main simulation code has been written in FORTRAN90. The code is available in the GitHub repository: https://github.com/PhyBi/Border-cell-migration/tree/main. The analyses of both the simulation and live imaging data have been carried out using our own MATLAB subroutines.

### MEASURED OBSERVABLES

#### Defining early and late phase of migration

We define the early 50% of migration as from the initiation of AP migration up to the time instant when the cluster has migrated 50% of the egg chamber length (L), and the rest of the migration up to the end of simulation, i.e., when the cluster has reached the oocyte boundary, is designated as the late 50%.

#### Calculating the position vector of the border cell cluster center

At an instant *t*, the position vector of a border cell center is defined as 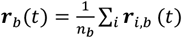. Similarly, the position vector of a polar cell center is defined as 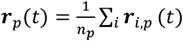. Here, *n*_*b*_ and *n*_*p*_ are the number of beads in each border and polar cell respectively. Now, the position vector of the cluster center at time instant *t* is ***R***_*b*_(*t*) = (∑_*i,b*_ ***r***_*i,b*_ (*t*) + ∑_*i,p*_ ***r***_*i,p*_ (*t*))/(*m*_*b*_*n*_*b*_ + *m*_*p*_*n*_*p*_). Here, *m*_*b*_ and *m*_*p*_ are the total number of border and polar cells considered in our system.

#### Net cluster speed in early and late phase

Suppose the ***R***_*b*_(*t*_0_), ***R***_*b*_(*t*_50_), ***R***_*b*_(*t*_100_) are the position vectors of the border cell cluster center at the time of initiation (*t*_0_) of AP migration, at 50% of migration (*t*_50_), and at completion of migration (*t*_100_) respectively. Then, net cluster speed in the early phase is defined as |***R***_*b*_(*t*_50_) − ***R***_*b*_(*t*_0_)|/(*t*_50_ − *t*_0_), and that in late phase is defined as |***R***_*b*_(*t*_100_) − ***R***_*b*_(*t*_50_)|/(*t*_100_ − *t*_50_).

#### AP migration and medial migration percentage

**Figure S2:**
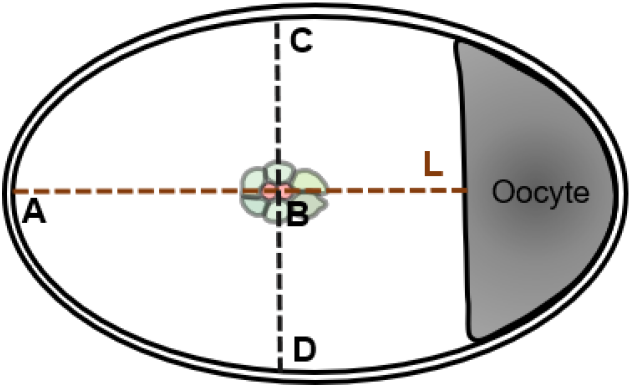
Measurement scheme of AP migration and medial migration percentages.

We measure the AP migration percentage of the cluster as the fraction of distance covered by the cluster center along the AP axis (i.e. the x-axis), defined as (AB/L)×100% (see Fig. S2) at the end of our observation time (500 simulation time units). Further at a time instant *t*, if the cluster center is at B (Fig. S2), we estimate the medial migration percentage defined as min(BD,BC)/(DC/2) ×100%. Higher medial migration percentage signifies the movement of the cluster along the center of the egg chamber, and lower medial migration implies a deviation from the central path.

#### Instantaneous cluster speed

Suppose the ***R***_*b*_(*t*), and ***R***_*b*_(*t* + Δ*t*) are the position vectors of the border cell cluster center at the two consecutive time instants that differ by Δ*t*, the smallest time resolution between two successive time frames. The instantaneous cluster speed at time instant *t* + Δ*t* is thus defined as |***R***_*b*_(*t* + Δ*t*) − ***R***_*b*_(*t*)|/Δ*t*.

#### Identifying positional switching and quantification scheme

At a time instant *t*, suppose ***r***_*b*_(*t*) denotes the position vector of a border cell nucleus and ***R***_*b*_(*t*) is the position vector of the border cell cluster center with respect to the origin. Now, to quantify the positional switching of the border cells, at every time instant we segregate the positions of the border cells within the cluster into three regions: front, side, and back, depending on the angle between the position vector of the border cell nucleus with respect to the cluster center (***r***_*b*_(*t*) − ***R***_*b*_(*t*)), and the instantaneous velocity vector of the cluster center ((***R***_*b*_(*t*) − ***R***_*b*_(*t* − Δ*t*))/Δ*t*) (Fig. 3C, main text). At an instant, if the magnitude of the angle is less than 45°, in between 45° to 135°, and greater than 135°, then the corresponding border cell is in the front, side, and back zone respectively. Finally, within the front zone, we further demarcate the leading border cell as the one having the smallest angle. Similarly, within the back zone, the border cell that has the largest angle, i.e. furthest from the instantaneous cluster velocity direction, gets identified as the trailing cell at the particular time instant. We register a single switching event if one of the leading or trailing cell identity or both get altered. A single switching event is denoted by 1 and an absence of any switching event is denoted by 0.

## TABLES

**Table S1.**
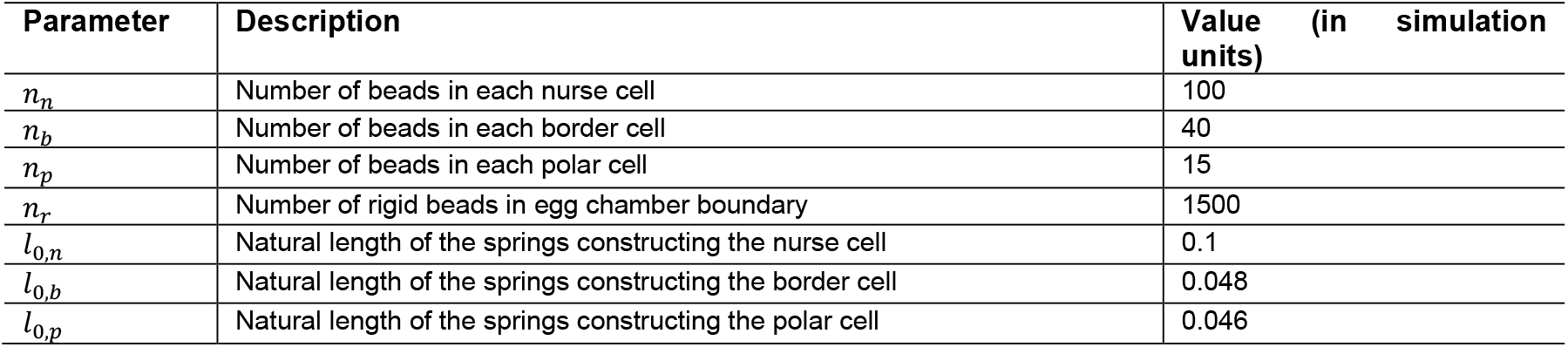
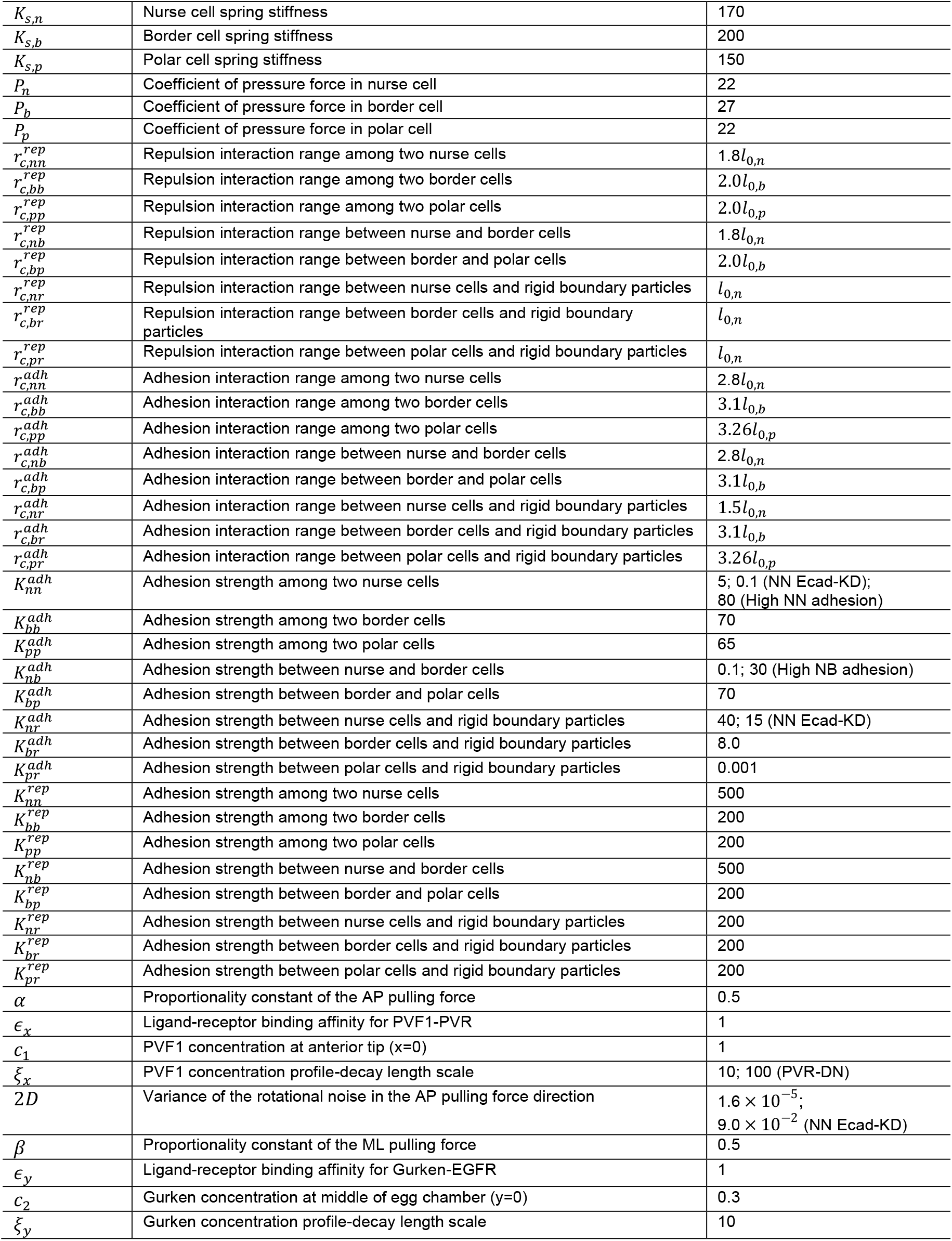
List of system parameters.

| Parameter | Description | Value (in simulation units) |
| --- | --- | --- |
| $n_n$ | Number of beads in each nurse cell | 100 |
| $n_b$ | Number of beads in each border cell | 40 |
| $n_p$ | Number of beads in each polar cell | 15 |
| $n_r$ | Number of rigid beads in egg chamber boundary | 1500 |
| $l_{0,n}$ | Natural length of the springs constructing the nurse cell | 0.1 |
| $l_{0,b}$ | Natural length of the springs constructing the border cell | 0.048 |
| $l_{0,p}$ | Natural length of the springs constructing the polar cell | 0.046 |
| $K_{s,n}$ | Nurse cell spring stiffness | 170 |
| $K_{s,b}$ | Border cell spring stiffness | 200 |
| $K_{s,p}$ | Polar cell spring stiffness | 150 |
| $P_n$ | Coefficient of pressure force in nurse cell | 22 |
| $P_b$ | Coefficient of pressure force in border cell | 27 |
| $P_p$ | Coefficient of pressure force in polar cell | 22 |
| $r_{c,nn}^{rep}$ | Repulsion interaction range among two nurse cells | $1.8l_{0,n}$ |
| $r_{c,bb}^{rep}$ | Repulsion interaction range among two border cells | $2.0l_{0,b}$ |
| $r_{c,pp}^{rep}$ | Repulsion interaction range among two polar cells | $2.0l_{0,p}$ |
| $r_{c,nb}^{rep}$ | Repulsion interaction range between nurse and border cells | $1.8l_{0,n}$ |
| $r_{c,bp}^{rep}$ | Repulsion interaction range between border and polar cells | $2.0l_{0,b}$ |
| $r_{c,nr}^{rep}$ | Repulsion interaction range between nurse cells and rigid boundary particles | $l_{0,n}$ |
| $r_{c,br}^{rep}$ | Repulsion interaction range between border cells and rigid boundary particles | $l_{0,n}$ |
| $r_{c,pr}^{rep}$ | Repulsion interaction range between polar cells and rigid boundary particles | $l_{0,n}$ |
| $r_{c,nn}^{adh}$ | Adhesion interaction range among two nurse cells | $2.8l_{0,n}$ |
| $r_{c,bb}^{adh}$ | Adhesion interaction range among two border cells | $3.1l_{0,b}$ |
| $r_{c,pp}^{adh}$ | Adhesion interaction range among two polar cells | $3.26l_{0,p}$ |
| $r_{c,nb}^{adh}$ | Adhesion interaction range between nurse and border cells | $2.8l_{0,n}$ |
| $r_{c,bp}^{adh}$ | Adhesion interaction range between border and polar cells | $3.1l_{0,b}$ |
| $r_{c,nr}^{adh}$ | Adhesion interaction range between nurse cells and rigid boundary particles | $1.5l_{0,n}$ |
| $r_{c,br}^{adh}$ | Adhesion interaction range between border cells and rigid boundary particles | $3.1l_{0,b}$ |
| $r_{c,pr}^{adh}$ | Adhesion interaction range between polar cells and rigid boundary particles | $3.26l_{0,p}$ |
| $K_{nn}^{adh}$ | Adhesion strength among two nurse cells | 5; 0.1 (NN Ecad-KD);<br>80 (High NN adhesion) |
| $K_{bb}^{adh}$ | Adhesion strength among two border cells | 70 |
| $K_{pp}^{adh}$ | Adhesion strength among two polar cells | 65 |
| $K_{nb}^{adh}$ | Adhesion strength between nurse and border cells | 0.1; 30 (High NB adhesion) |
| $K_{bp}^{adh}$ | Adhesion strength between border and polar cells | 70 |
| $K_{nr}^{adh}$ | Adhesion strength between nurse cells and rigid boundary particles | 40; 15 (NN Ecad-KD) |
| $K_{br}^{adh}$ | Adhesion strength between border cells and rigid boundary particles | 8.0 |
| $K_{pr}^{adh}$ | Adhesion strength between polar cells and rigid boundary particles | 0.001 |
| $K_{nn}^{rep}$ | Adhesion strength among two nurse cells | 500 |
| $K_{bb}^{rep}$ | Adhesion strength among two border cells | 200 |
| $K_{pp}^{rep}$ | Adhesion strength among two polar cells | 200 |
| $K_{nb}^{rep}$ | Adhesion strength between nurse and border cells | 500 |
| $K_{bp}^{rep}$ | Adhesion strength between border and polar cells | 200 |
| $K_{nr}^{rep}$ | Adhesion strength between nurse cells and rigid boundary particles | 200 |
| $K_{br}^{rep}$ | Adhesion strength between border cells and rigid boundary particles | 200 |
| $K_{pr}^{rep}$ | Adhesion strength between polar cells and rigid boundary particles | 200 |
| $\alpha$ | Proportionality constant of the AP pulling force | 0.5 |
| $\epsilon_x$ | Ligand-receptor binding affinity for PVF1-PVR | 1 |
| $c_1$ | PVF1 concentration at anterior tip ($x=0$) | 1 |
| $\xi_x$ | PVF1 concentration profile-decay length scale | 10; 100 (PVR-DN) |
| $2D$ | Variance of the rotational noise in the AP pulling force direction | $1.6 \times 10^{-5}$;<br>$9.0 \times 10^{-2}$ (NN Ecad-KD) |
| $\beta$ | Proportionality constant of the ML pulling force | 0.5 |
| $\epsilon_y$ | Ligand-receptor binding affinity for Gurken-EGFR | 1 |
| $c_2$ | Gurken concentration at middle of egg chamber ($y=0$) | 0.3 |
| $\xi_y$ | Gurken concentration profile-decay length scale | 10 |

## MOVIE CAPTIONS

**Movie S1. Control time-lapse confocal movie of border cell migration in *Drosophila* egg chamber of genotype**, *slbo*-Gal4,UAS-mCD8:GFP; UAS-*stinger-NLS*. The border cell membranes are marked in green (GFP), the nuclei of border and polar cells are marked in red (Stinger-NLS) and grey is the egg chamber captured with DIC filter. Colored trajectories show the path of the tracked border cell nuclei through the course of migration, where the color gradient resembles migration time. Notably, near a multi-nurse cells junction, from frame number 17-20, the cluster substantially speeds up and then the speed falls down, indicating the acceleration and deceleration phases of migration.

**Movie S2. A 2D wild-type simulation video showing the full AP migration of the border cell cluster**. The larger grey colored cells are the nurse cells, green and red colored cells are the border and the polar cells respectively. Note that the cluster accelerates near the multi-nurse cells junctions and decelerates while it unzips a two-cell junction.

**Movie S3. A wild-type simulation video showing the cluster rotation in 2D**. The cluster rotates near a three-nurse cells junction.

**Movie S4. A simulation video in nurse-nurse E-cadherin KD condition showing a disrupted medial migration**.

